# High-quality complete genome resource of plant pathogenic bacterium *Dickeya solani* strain IPO 2019 isolated from *Hyacinthus orientalis*

**DOI:** 10.1101/2021.03.02.433526

**Authors:** Robert Czajkowski, Lukasz Rabalski, Przemysław Bartnik, Sylwia Jafra

## Abstract

*Dickeya solani* is an emerging plant pathogenic bacterium, causing disease symptoms in a variety of agriculturally relevant crop species worldwide. To date a number of *D. solani* genomes have been sequenced and characterized, the great majority of these genomes have however come from *D. solani* strains isolated from potato (*Solanum tuberosum* L.) and not from other plant hosts. Herewith, we present the first complete, high-quality genome of *D. solani* strain IPO 2019 (LMG 25990) isolated from ornamental plant *Hyacinthus orientalis*. The genome of *D. solani* strain IPO 2019 consists of one chromosome of 4,919,542 bp., with a GC content of 56.2% and no plasmids. The genome contains 4502 annotated features, 22 rRNA genes, 73 tRNA genes and 1 CRISPRS. We believe that the information of this high-quality, complete, closed genome of *D. solani* strain isolated from host plant different than potato (i.e. hyacinth) will provide resources for comparative genomic studies as well as for analyses targeting adaptation and ecological fitness mechanisms present in *Dickeya solani* species.

## Genome announcement

*Dickeya solani* (former *Dickeya* spp. biovar 3) is an emerging plant pathogenic bacterium, causing disease symptoms in a variety of agricultural plant species worldwide (Toth et al., 2011; van der Wolf et al., 2014). The pathogen was first reported in potato in early 2000s as a distinct genetic clade of biovar 3, causing severe outbreaks in potato cultivation in different European countries (Czajkowski et al., 2009; Slawiak et al., 2009; Laurila et al., 2010). Until now the pathogen has been repeatably isolated from symptomatic as well as from symptomless potato plants grown in agricultural fields across Europe and worldwide (van der Wolf et al., 2014). As *D. solani* was reported to cause severe disease symptoms under hot climatic conditions as well as there is an observed relationship between *D. solani* high disease incidences in crops and the presence of hot growing seasons in temperate countries (Degefu et al., 2013; Tsror et al., 2013) it has been speculated that the bacterium has been introduced from subtropical and/or tropical agricultural regions outside Europe to Europe (Slawiak et al., 2009; Toth et al., 2011). Likewise, it has been suggested that *D. solani* might have first appeared in ornamental plants cultivated commercially outside Europe and/or in plants grown in Europe in glasshouses under elevated temperature regime and from there spread to potato ecosystem in temperate regions (Slawiak et al., 2009). At the moment, due to the limited number of *D. solani* strains isolated from different (e.g. (sub)tropical, ornamental) plant hosts, there are not enough data supporting either of these hypotheses. It is well accepted however that the pathogen has been only recently introduced to potato ecosystem, as most strains of *D. solani* analyzed belong to the same haplotype and expresses only a low number of genetic variations (Khayi et al., 2016).

Since the establishment of *D. solani* species in 2014 (van der Wolf et al., 2014), the strain IPO 2222 (DSM 28711, LMG 25993, NCPPB 4479) is recognized as a type strain. A complete, closed, high quality genome sequence of *D. solani* strain IPO 2222 isolated from potato together with the corresponding phylogenetic and phenotypic information about the IPO 2222 has been published (Khayi et al., 2016). Likewise, to date, a number of *D. solani* genomes have been sequenced and characterized (https://www.ncbi.nlm.nih.gov/genome/14829). At the moment of manuscript preparation (February 2021) the great majority of these (completed and draft) genome sequences however have come from *D. solani* strains isolated exclusively from potato and not from other plant hosts.

*D. solani* strain IPO 2019 was isolated from *Hyacinthus orientalis* (common name: garden hyacinth, Dutch hyacinth) in 2009 in the Netherlands. The strain has been deposited in the Belgian Co-ordinated Collection of Micro-organisms (BCCM) by Johan van Vaerenbergh (ILVO Eenheid Plant, Onderzoeksdomein Gewasbescherming, Belgium) under collection number LMG 25990. In our comparative phenotypic studies involving *D. solani* strains IPO 2019 and IPO 2222, we detected that IPO 2019 but not IPO 2222 was able to utilize a unique panel of saccharides including dextrin, D-maltose, D-trehalose, gentobiose, D-turanose, L-rhamnose and sorbitol as carbon sources (Bartnik, Czajkowski, unpublished). To further explore the hypothesis that IPO 2019 isolated from hyacinth may show distinct phenotypic and genetic characteristics from *D. solani* type strain IPO 2222 isolated from potato and to link the preliminarily observed phenotypes with the genomic data, the genome of IPO 2019 strain has been sequenced and annotated.

For isolation of genomic DNA, *D. solani* IPO 2019 was grown at 28 °C in tryptone soya broth (TSB, Oxoid) for 16 h with shaking (150 rpm). Bacterial DNA was isolated using a Wizard Genomic DNA Purification Kit (Promega) according to the protocol provided by the manufacturer. The IPO 2019 genome was sequenced by a hybrid approach with long reads generated using Oxford Nanopore Technologies (ONT) and short reads generated using Illumina technology. DNA sequencing libraries were prepared according to manufacturer (for ONT it was LSK kit and for Illumina it was Nextera XT kit). Long reads were generated by single Flongle run on MinION and short reads were generated during Midoutput run on Ilumina Miniseq. *De novo* assembly of long reads was performed using Flye 2.8.2. (https://github.com/fenderglass/Flye) with final mean coverage × 28. Initial genome polishing was conducted by tool integrated in the assembler. Pilon 1.23 (https://github.com/broadinstitute/pilon) was used to further error corrections. Final mean coverage of short reds to polished genome was 748.9. The combined procedure produced a single contig of 4,919,542 bp., which was further manually inspected and curated. Repetitive genetic elements were searched using Geneious (Kearse et al., 2012). Finally, BLAST (http://blast.ncbi.nlm.nih.gov/Blast.cgi), InterProScan (http://www.ebi.ac.uk/Tools/pfa/iprscan/) and HMMER (phmmer, UniProtKB) (Finn et al., 2011) (http://hmmer.org/) were used to annotate the IPO 2019 genome. The obtained structural and functional annotations were retested using RAST (Rapid Annotation using Subsystem Technology (http://rast.nmpdr.org/) (Aziz et al., 2008) and DFAST (http://dfast.ddbj.nig.ac.jp/) (Tanizawa et al., 2018). Preliminary comparative genomics analyses (IPO 2019 vs. IPO 2222) were done using EDGAR (Blom et al., 2009; Blom et al., 2016). The presence of prophages (bacteriophage sequences) in the genome of IPO 2019 was predicted using PHASTER (Arndt et al., 2016) and the presence of gene clusters involved in biosynthesis of secondary metabolites was predicted using anitSMASH 5.0 database (Blin et al., 2019).

The complete, closed, high-quality genome of *D. solani* strain IPO 2019 has been uploaded to NCBI Genbank and received the accession number: CP071062. Upon submission to NCBI Genbank, the genome was also annotated by the NCBI Prokaryotic Genome Annotation Pipeline (https://www.ncbi.nlm.nih.gov/genome/annotation_prok/) (Tatusova et al., 2016). The genome is double-stranded 4,919,542 bp. DNA sequence, possessing 4502 annotated features and an average GC content of 56.2% with no plasmids. The average gene length was predicted to be ca. 1200 bp. and the average protein length was predicted to be ca. 333 amino acids. 85.8% of the genome consists of coding regions. The *D. solani* IPO 2019 genome encodes 22 rRNAs, 73 tRNAs and 1 CRISPRS. We also identified 11 gene clusters responsible for synthesis of secondary metabolites. Six of these matched known clusters for the biosynthesis of NRPS (non-ribosomal peptide synthases), batalactone, cyanobacin, arylpolyene and thiopeptide. The investigation of IPO 2019 genome for presence of viral sequences done with PHASTER did not result in prediction of regions containing intact (complete) viral genomes, three regions in the genome were however found containing incomplete prophage genomes.

The first, completed, high-quality genome of *D. solani* strain isolated from hyacinth plant may prove useful in (comparative) adaptation studies and ecological fitness analyses done to better understand the origin, spread and virulence traits of this new, emerging *Dickeya* species.

## Authors’ statement

The authors declare than no conflict of interest exists.

## Acknowledgements

The authors would like to thank Dr. Jan M. van der Wolf (Plant Research International, Wageningen University and Research Center, Wageningen, The Netherlands) for providing *D. solani* strain IPO 2019 for this study. The work was financially supported by Polish Ministry of Higher Education (Ministers two Nauki i Szkolnictwa Wyz□szego, Polska) funds DS 531-N104-D800-21 to RC and by Polish Ministry of Science and Higher Education (Ministerstwo Nauki i Szkolnictwa Wyz□szego, Polska) funds DS 531-M105-D786-21 to SJ.

---

**Figure 1.**
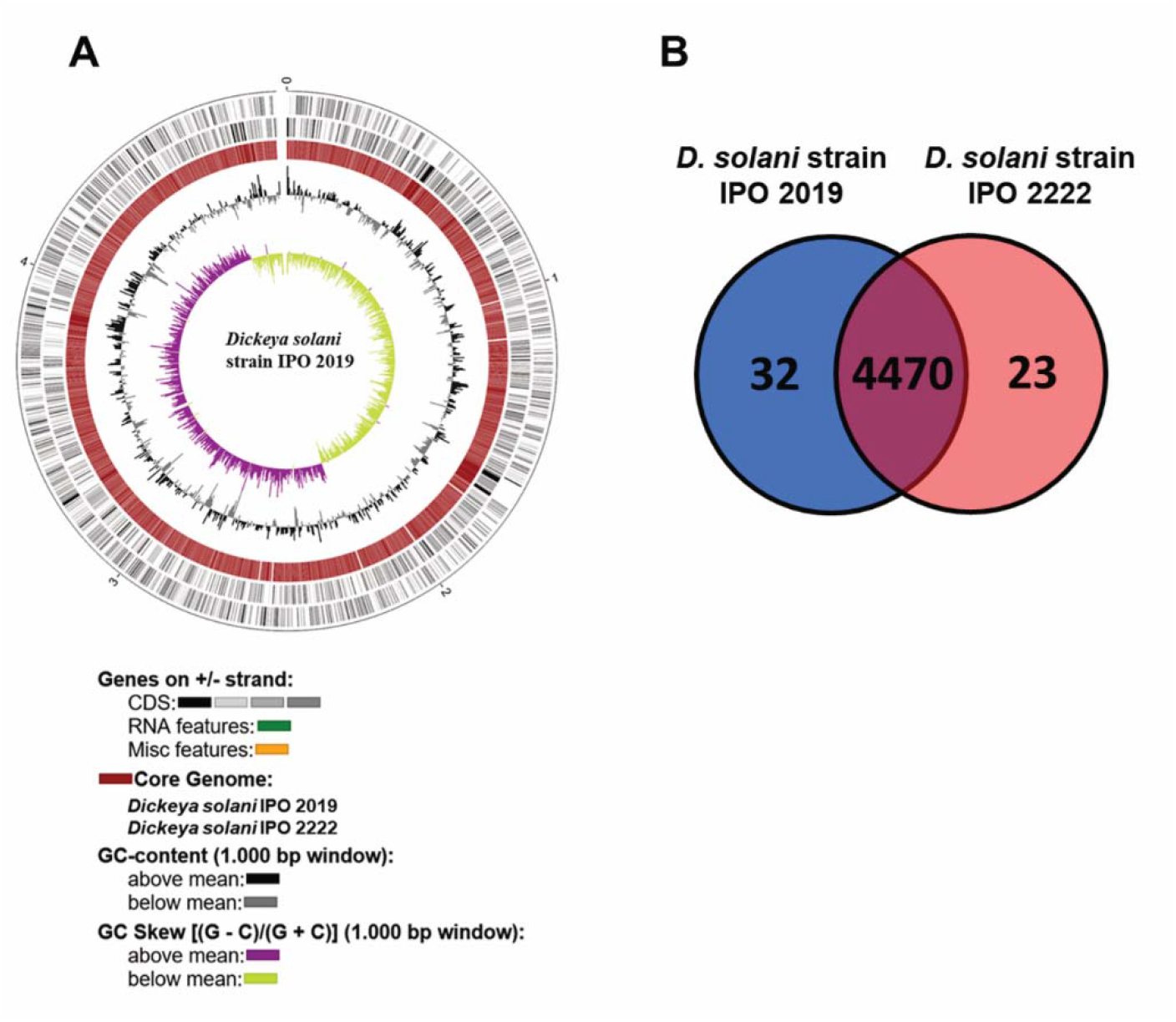
Features of the *D. solani* strain IPO 2019 genome and its comparison with the IPO 2222 genome. Circular plot of the IPO 2019 and IPO 2222 genomes (**A**) and core genome shared between D. solani strains IPO 2019 and IPO 2222 (**B**). The comparative analyses were done using EDGAR – an enhanced software platform for comparative gene content analyses (Blom et al., 2016).

**Table 1.** General features of the *D. solani* strain IPO 2019 complete genome.

|  | **IPO 2019 genome** |
| --- | --- |
| **Genome size (bp.)** | 4,919,542 |
| **GC content (%)** | 56.2 |
| **No. contigs** | 1 |
| **N50** | 1 |
| **Coding ratio (%)** | 85.8 |
| **No. total coding sequences (features)** | 4502 |
| **Average gene length (bp.)** | 1200 |
| **Presence of plasmids** | no |
| **rRNAs** | 22 |
| **tRNAs** | 73 |
| **CRISPRs** | 1 |
| **No. intact prophage genomes**<br>**(according to PHASTER)** | 0 |
| **GenBank accession number** | CP071062 |

